# The Arabidopsis *SNARE* complex genes regulate the early stages of pollen-stigma interactions

**DOI:** 10.1101/2023.10.16.562513

**Authors:** Stuart R. Macgregor, Paula K.S. Beronilla, Daphne R. Goring

## Abstract

In the Brassicaceae, the process of accepting compatible pollen is a key step in successful reproduction and highly regulated following interactions between the pollen and the stigma. Central to this, is the initiation of secretion in the stigma, which is proposed to provide resources to the pollen for hydration and germination and pollen tube growth. Previously, the eight exocyst subunit genes were shown to be required in the Arabidopsis stigma to support these pollen responses. One of the roles of the exocyst is to tether secretory vesicles at the plasma membrane for membrane fusion by the SNARE complex to enable vesicle cargo release. Here, we investigate the role of Arabidopsis *SNARE* genes in the stigma for pollen responses. Using a combination of different knockout and knockdown *SNARE* mutant lines, we show that *VAMP721, VAMP722, SYP121, SYP122* and *SNAP33* are involved in this process. Significant disruptions in pollen hydration were observed following pollination of wildtype pollen on the mutant *SNARE* stigmas. Overall, these results place the Arabidopsis SNARE complex as a contributor in the stigma for pollen responses and reaffirm the significance of secretion in the stigma to support the pollen-stigma interactions.

**Key Message:** The VAMP721, VAMP722, SYP121, SYP122 and SNAP33 SNAREs are required in the Arabidopsis stigma for pollen hydration, further supporting a role for vesicle trafficking in the stigma’s pollen responses.

## Introduction

Following pollination in *Arabidopsis thaliana* (Arabidopsis), the first point of pollen grain contact occurs with the stigma at the top of the pistil. The desiccated pollen grain is reliant on the stigmatic papillae on the surface of the stigma to provide water for hydration, and this occurs when the pollen is recognized as compatible (which is self-pollen in this naturally-selfing species). Following hydration, the pollen grain germinates to form a pollen tube that enters the stigma and grows down the reproductive tract to an ovule where sperm cells are released for gamete fusion, initiating seed development (Hafidh and Honys 2021; Kim et al. 2021; Robichaux and Wallace 2021). The pollen hydration stage is an important initial control point in this process (Rozier et al. 2020), and the cellular pathways in the stigmatic papillae regulating compatible pollen recognition and responses are still not fully understood. The pollen grain has a surface pollen coat containing a number of different small cysteine-rich proteins (CRPs) which could act as ligands for receptors in the stigma papillae (Wang et al. 2023), and one set of Arabidopsis CRPs, the Pollen Coat Protein Bs (PCP-Bs), are required for full pollen hydration (Wang et al. 2017). The Arabidopsis FERONIA and ANJEA receptor kinases are proposed to act in the stigma as negative regulators of pollen hydration by maintaining high levels of intracellular ROS in the stigmatic papillae. Following pollination, PCP-Bs are proposed to bind and inhibit these receptor kinases to decrease ROS levels thereby allowing pollen hydration (Liu et al. 2021). Another group of Arabidopsis receptor-like kinase genes, RKF1 and paralogues, are proposed to function in the stigma as positive regulators of pollen hydration (Bordeleau et al. 2022; Lee and Goring 2021).

One of the cellular responses in the stigma linked to pollen hydration involves vesicle trafficking in the stigmatic papilla to presumably deliver cargo that aids in water release to the pollen grain (Goring 2017; Safavian and Goring 2013). Within the secretory pathway, the final stages of vesicle delivery to the PM has been extensively investigated in eukaryotes, and key protein complexes have been identified to contribute to this final stage of secretion, including a tethering complex known as the exocyst complex followed by the SOLUBLE N-ETHYLMALEIMIDE-SENSITIVE FACTOR ADAPTOR PROTEIN RECEPTOR (SNARE) complex (Lipka et al. 2007; Saeed et al. 2019). Both protein complexes are essential for a large diversity of secretory events throughout the plant lifecycle, and corresponding mutants often display altered development and seedling lethality (Luo et al. 2022; Shi et al. 2023; Zarsky 2022; Zarsky et al. 2020). The octameric exocyst complex is involved in tethering post-Golgi secretory vesicles at the plasma membrane (Saeed et al. 2019; Zarsky 2022). Disrupting the expression of the corresponding Arabidopsis exocyst subunit genes for EXO70A1, SEC3, SEC5, SEC6, SEC8, SEC10, SEC15, and EXO84 resulted in a significant reduction in wildtype pollen hydration and pollen germination on the mutant stigmas indicating that the exocyst complex was essential for pollen responses in the Arabidopsis stigma (Safavian et al. 2014; Safavian et al. 2015; Samuel et al. 2009). In addition, a MAPK pathway regulates EXO70A1 to confer stigma receptivity (Jamshed et al. 2020). Interestingly, one of the components, EXO70A1, is also targeted by the self-incompatibility pathway in the stigma to reject self-incompatible pollen (Abhinandan et al. 2022; Samuel et al. 2009; Sankaranarayanan et al. 2015). Vesicle docking by the exocyst complex at the plasma membrane is typically followed by membrane fusion facilitated by the SNARE complex (Luo et al. 2022; Vukasinovic and Zarsky 2016). The SNARE complex carries out this function through protein-protein interactions between R-SNAREs on the vesicle membrane and Q-SNAREs on the target membrane (e.g. plasma membrane) to form a fusion complex for membrane fusion and release of the vesicle content (Lipka et al. 2007; Shi et al. 2023).

With the exocyst complex implicated in the stigma for supporting pollen hydration and germination, we next were interested in investigating the SNARE genes that could also be supporting this process. The Arabidopsis genome contains 64 predicted SNARE genes, defined as those containing a SNARE domain that mediates the protein-protein interactions required for final SNARE complex assembly. Many of these SNARE proteins are involved in other stages of vesicle trafficking in the endomembrane system (e.g. ER/Golgi, TGN/endosome, vacuole) with 23 SNARE proteins associated with vesicle fusion at the plasma membrane (Luo et al. 2022; Shi et al. 2023). As with many other Arabidopsis gene families, the SNARE gene family has single members that are critical for plant growth as well as closely related family members with overlapping roles. For this study, we focused on two R-SNARE genes, *VESICLE ASSOCIATED MEMBRANE PROTEIN* (*VAMP*)*721* and *VAMP722*, and three plasma membrane-associated Q-SNARE genes, *SYNTAXIN IN PLANTS* (*SYP*)*121, SYP122* and *SOLUBLE N-ETHYLMALEIMIDE-SENSITIVE FACTOR ADAPTOR PROTEIN* (*SNAP*)*33*, as based on previous work, they represented strong candidates for regulating pollen-stigma interactions. All five SNAREs have been implicated in working together in Arabidopsis immune responses against powdery mildew infections (Kwon et al. 2008; Rubiato et al. 2022; Yun et al. 2023). As well, SYP121 was found to form a protein complex with SNAP33 and VAMP721/722 in response to powdery mildew infection (Kwon et al. 2008). SNAREs function by forming fusion complexes for membrane fusion, and one type of complex consists of a vesicle R-SNARE (VAMP721/722) with the plasma membrane Qa-SNARE (SYP121/122) and Qbc-SNARE (SNAP33) (Lipka et al. 2007). A connection between the exocyst complex and these SNAREs was demonstrated with EXO70A1 shown to interact with SYP121, SNAP33 and VAMP721/722 (Larson et al. 2020). As well, decreased VAMP721 trafficking to the plasma membrane was observed in the *exo70a1*mutant (Fendrych et al. 2013). Finally, SYP121 has been implicated in trafficking the PIP2:7 aquaporin to the plasma membrane (Hachez et al. 2014; Laloux et al. 2020), and more recently a role for aquaporins in facilitating pollen hydration has been reported (Windari et al. 2021).

One of the challenges in working with mutants for *VAMP721, VAMP722, SYP121, SYP122* and *SNAP33* is that these *SNARE*s have also been implicated in secretory pathways regulating Arabidopsis growth and development, and loss-of-function mutations lead to impaired growth and seedling lethality (Luo et al. 2022; Shi et al. 2023). The single *vamp721* and *vamp722* mutants are wildtype in appearance, but the *vamp721 vamp722* double mutant displays a seedling-lethal phenotype with defects in cell plate formation during cytokinesis (Kwon et al. 2008; Zhang et al. 2011). The homozygous *snap33* mutant also displays early seedling death with mild cell plate formation defects and a lesion-mimic phenotype causing necrotic lesions on the leaves (El Kasmi et al. 2013; Heese et al. 2001; Henchiri 2021). Finally, the single *syp121* and *syp122* mutants display wildtype plant growth, but the *syp121 syp122* double mutant is a severely dwarfed lesion-mimic mutant with highly elevated levels of salicylic acid (Zhang et al. 2007; Zhang et al. 2008). These phenotypes in the *syp121 syp122* double mutant can be rescued by expressing the bacterial *NahG* transgene which encodes a salicylate hydroxylase to metabolize and reduce SA levels in the plant (Gaffney et al. 1993; Zhang et al. 2008). Thus, to study the role of these five *SNARE* genes in the early stages of pollen-stigma interactions, we adopted knockout/knockdown strategies to reduce the expression of these genes to the lowest possible level without causing lethality.

## Material and methods

### Arabidopsis SNARE mutants and growth conditions

The five SNARE genes selected for this study were *VAMP721* (At1g04750), *VAMP722* (At2g33120), *SYP121* (At3g11820), *SYP122* (At3g52400) and *SNAP33* (At5g61210). Expression profiles for these genes were retrieved from public RNA-Seq datasets (Gao et al. 2018; Klepikova et al. 2016) and heatmaps were generated using the BAR HeatMapper Plus Tool (Toufighi et al. 2005); Supplementary Fig. 1). The following previously characterized T-DNA mutants in the Col-0 background were used (Supplementary Fig. 2): *vamp721-1* (SALK_037273; (Kwon et al. 2008), *vamp722-1* (SALK_119149; (Kwon et al. 2008) *syp122-1* (SALK_008617; (Assaad et al. 2004) and *snap33-2* (SALK_063806; (Henchiri 2021). For *SYP121*, the *syp121-5* mutant in the Col-0 background was created using CRISR/Cas9 (Supplementary Fig. 3). All mutants were confirmed by PCR genotyping using primers listed in Supplementary Table 1. All Arabidopsis seeds were cold stratified for at least 2 days at 4°C, then sowed directly onto soil (Sunshine #1). All soil was supplemented with 1g/L of 20-20-20 fertilizer and grown at 22°C on a 16-hour light/8-hour dark cycle. Humidity was monitored in both the growth chamber and laboratory and maintained between 20 and 60% relative humidity.

**Table I.**
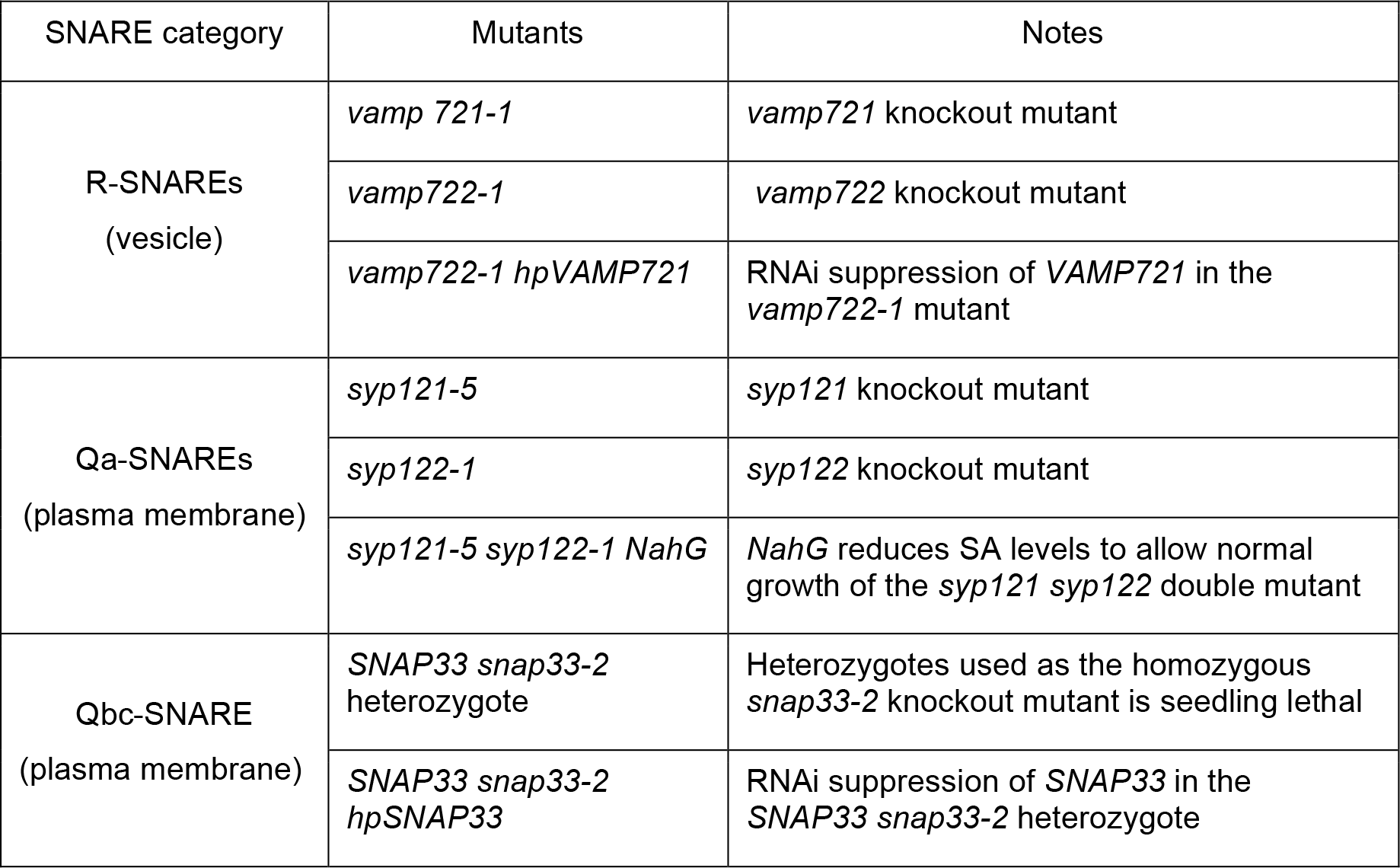
Mutants used in this study.

| SNARE category | Mutants | Notes |
| --- | --- | --- |
| R-SNAREs<br>(vesicle) | *vamp 721-1* | *vamp721* knockout mutant |
|  | *vamp722-1* | *vamp722* knockout mutant |
|  | *vamp722-1 hpVAMP721* | RNAi suppression of *VAMP721* in the *vamp722-1* mutant |
| Qa-SNAREs<br>(plasma membrane) | *syp121-5* | *syp121* knockout mutant |
|  | *syp122-1* | *syp122* knockout mutant |
|  | *syp121-5 syp122-1 NahG* | *NahG* reduces SA levels to allow normal growth of the *syp121 syp122* double mutant |
| Qbc-SNARE<br>(plasma membrane) | *SNAP33 snap33-2* heterozygote | Heterozygotes used as the homozygous *snap33-2* knockout mutant is seedling lethal |
|  | *SNAP33 snap33-2 hpSNAP33* | RNAi suppression of *SNAP33* in the *SNAP33 snap33-2* heterozygote |

### Plasmid construction and plant transformation

The *syp121-5* CRISPR/Cas9 deletion mutant was generated in the Col-0 background using previously described methods (Doucet et al. 2019; Wang et al. 2015). The final pBEE401E vector contained two guide RNAs targeting exon 1 and 2 of *SYP121* (Supplementary Fig. 3). Arabidopsis Col-0 plants were transformed using floral dip (Clough and Bent, 1998). T1 seeds were collected and germinated on soil, and one-week old seedlings were sprayed with BASTA herbicide to select for transformants. PCR was used to confirm transformants (BASTA resistance marker), and the presence of *syp121* mutations (deletions of the exon 1-2 region). An additional round of PCR was used to confirm heterozygosity of the deletions using primers flanking the guide RNA target sites. As shown for other *syp121* mutant alleles, *syp121-5* is morphologically wildtype in development and floral morphology, but displays a lesion mimic phenotype when crossed with *syp122-1* (Supplementary Fig. 3). Homozygous T3 plants were used in the crosses between *syp121-5* and *syp122-1*, and plants obtained from this cross (*syp121-5* SYP122 *syp122-1* or SYP121 *syp121-5 syp122*-1) were then crossed with the *NahG* transgenic Col-0 plants (Shah et al. 1999). The resulting F_2_ double homozygous (*syp121-5 syp122-1 NahG*) plants were used for all experiments.

### RNA-silencing plasmid constructs

Hairpin RNA-induced silencing (hpRNAi) plasmids under the control of the stigma-specific SLR1 promoter and targeting *SNAP33* or *VAMP721* were constructed using gateway cloning. First, the Gateway hairpin fragment from pB7GWIWG2(II) gateway vector (Karimi et al. 2002) was synthesized by GeneArt (ThermoFisher) and cloned into pORE3:SLR1p using the added ClaI and SacI restriction enzyme sites resulting in the pORE3-SLR1::GWhpCassette plasmid. cDNA fragments from *SNAP33* or *VAMP721* (Supplementary Fig. 2) were gateway cloned into SLR1::GWhpCassette plasmid to generate the hpRNAi plasmids which were then transformed into their respective T-DNA mutants via *Agrobacterium* mediated floral dipping.

### Assays for Pollen Hydration, Aniline Blue Staining and Seed set

All post-pollination assays were performed with manual pollinations and conducted as described in (Lee et al. 2020). For all assays, late stage 12 pistils were first emasculated and wrapped in plastic wrap overnight prior to pollination. To ensure consistency, all pollinations were performed at humidity levels below 50%. The pollen hydration assay was carried out by removing and mounting pistils on ½ MS medium, lightly pollinating the stigmatic surface with a single Col-0 anther and taking images at 0 and 10 minutes after pollination using a Nikon SMZ800 stereo zoom microscope system. Randomly selected pollen grains were measured using the NIS-elements imaging software. For each timepoint, 30 pollen grains were measured (3 stigmas with 10 pollen grains/stigma) and the experiment was replicated as least 2 times. For aniline blue staining, one Col-0 anther was used to gently pollinate each pistil, after which the plant was returned to a growth chamber at below 50% humidity. The pistils were collected at 2-hours or 24-hours post-pollination for staining, as described in Lee et al. (2020). The aniline blue-stained pistils were then mounted on a slide with sterile water, and imaged using a Zeiss Axioskop 2 Plus fluorescence microscope. There was a minimum of 8 pollinated pistils stained per each line. For seed counts, pistils from previously emasculated flowers were manually pollinated using a single Col-0 anther and left to develop in a growth chamber for 14 days (n=10 siliques/line). Once matured, the siliques were collected and cleared in 70% ethanol (v/v) for 5 days, or until seeds were clearly visible (Beuder et al. 2020). The cleared siliques were then mounted on a dry slide, with the septum facing upwards, and imaged using the Nikon sMz800 microscope. Seeds in the developed siliques were counted using the NIS-elements imaging software. To count ovules, mature ovules from naturally pollinated siliques were counted as described in (Yuan and Kessler 2019).

## Results

The aim of this study was to investigate the function of Arabidopsis *R-, Qa-* and *Qbc-SNARE* genes in the stigma to support early pollen-stigma interactions. We focused on the vesicle-associated *VAMP721* and *VAMP722* R-SNAREs, and the plasma membrane-associated *SYP121* and *SYP122* Qa-SNAREs and the *SNAP33* Qbc-SNARE. As part of the process in selecting these five *SNARE* genes, we investigated the expression profiles of *SNARE* genes with functions associated at the plasma membrane (Luo et al. 2022) in RNA-seq datasets covering a wide range of tissues including stigmas and stigmatic papillae (Gao et al. 2018; Klepikova et al. 2016). *VAMP721* and *VAMP722* are the most highly expressed *R-SNAREs* in the stigma and the stigmatic papillae datasets (Supplementary Fig. 1). Since they can function redundantly, the goal was to examine both the single and double mutants. However, while the single *vamp721-1* and *vamp722-1* mutants are wildtype in appearance, the *vamp721 vamp722* double mutant displays a seedling-lethal phenotype (Kwon et al. 2008; Zhang et al. 2011). As an alternative to the double mutant, an hpRNAi construct targeting *VAMP721* (*hpVAMP721*; Supplementary Fig. 2) under the control of the stigma-specific SLR1 promoter was transformed into the *vamp722-1* mutant. The *vamp722-1 hpVAMP721* plants displayed wildtype growth and flowering. Expression analysis of *VAMP721* via qPCR using RNA collected from half-pistil tissue indicated a significant reduction in *VAMP721* RNA levels when compared to Col-0 pistils (Supplementary Fig. 4).

For the *Qa-SNARE* genes, *SYP132* and *SYP121* displayed the highest expression in the stigma and the stigmatic papillae datasets (Supplementary Fig. 1). However, we ended up selecting *SYP121* and *SYP122* for this analysis since the two genes can function redundantly (Zhang et al. 2007; Zhang et al. 2008), and the proteins were found to form a complex with VAMP721, VAMP722 and SNAP33 (Kwon et al. 2008). For *SYP121*, CRISPR/Cas9 was used to generate the *syp121-5* deletion mutant to track in crosses more easily (Supplementary Fig. 3). The *syp121-5* mutant and the *syp122-1* T-DNA mutant (Assaad et al. 2004) are wildtype in appearance. As the double mutant displays growth defects, *syp121-5* and *syp122-1* were crossed with each other and to the *NahG* transgenic line (Shah et al. 1999) to generate the *syp122-1 syp121-5* double mutant with *NahG* which rescued the growth defects associated with elevated SA levels in the double mutant (Zhang et al. 2008); Supplementary Fig. 3).

For the *Qbc-SNARE* genes, *SNAP33* is the only member significantly expressed in the stigma and the stigmatic papillae datasets tissue (Supplementary Fig. 1). The homozygous *snap33* displays a seedling lethal phenotype (Heese et al. 2001) and so data was collected on heterozygous *SNAP33/snap33-2* plants. In addition, an hpRNAi construct targeting *SNAP33* (*hp SNAP33*; Supplementary Fig. 2) under the control of the SLR1 promoter was transformed into the *SNAP33/snap33-2* heterozygote. Data was collected on *SNAP33/snap33-2 hp SNAP33* plants which displayed wildtype growth and flowering. Expression analysis of *SNAP33* via qPCR indicated a significant reduction in *SNAP33* RNA levels in the *SNAP33/snap33-2 hpSNAP33* half-pistil samples (Supplementary Fig. 5).

### Wildtype Col-0 pollen hydration on *SNARE* mutant stigmas

One of the earliest checkpoints following pollination in Arabidopsis is the rapid hydration of pollen within the first 10 minutes of pollen-stigmatic papilla contact. Previously, we had found that mutant stigmas for different exocyst subunits supported reduced hydration of wildtype pollen at 10 min post-pollination (Safavian et al. 2015). Therefore, we used this same assay to measure how well the *SNARE* mutant stigmas supported wildtype Col-0 pollen hydration. In this assay, the Col-0 pollen grain becomes rounder as it hydrates, and the pollen diameter is measured at 0 and 10 min as a proxy for hydration (Lee et al. 2020). When Col-0 pollen was placed on stigmas from either of the single *vamp721-1* or *vamp722*-1 mutant stigmas, there was a significant decrease in pollen hydration at 10 minutes compared to Col-0 implicating both of these R-SNAREs in this process (Fig. 1c). Furthermore, when the same Col-0 pollen hydration assay was conducted on the three *vamp722-1 hpVAMP721* transgenic lines, there was a further significant reduction in the degree of Col-0 pollen hydration on these mutant stigmas at 10 minutes post-pollination (Fig 1a-c). This indicated that the reduced *VAMP721* expression in the *vamp722-1* mutant stigmas further impaired the ability of these stigmas to support Col-0 pollen hydration. For the *syp121-5* and *syp122-1* mutants, Col-0 pollen placed on the single mutant stigmas also displayed reduced hydration at 10 minutes. Interestingly, Col-0 pollen hydration on stigmas from the double *syp121-5 syp122-1* mutant expressing *NahG* displayed the same level of reduced hydration as the single mutants (Fig. 1d) rather than the additive effect that was seen for the *vamp722-1 hpVAMP721* transgenic lines. Together, these data indicate that *SYP121* and *SYP122* are likely playing some role in the stigma facilitating Col-0 pollen hydration. Finally, this assay also implicated the *SNAP33 Qbc-SNARE* gene as Col-0 pollen hydration assays on stigmas from the *SNAP33 snap33-2* heterozygote displayed a decrease in Col-0 pollen hydration at 10 minutes. When *SNAP33* expression levels were further decreased in the *SNAP33 snap33-2 hpSNAP33* stigmas, there was a further reduction in Col-0 pollen hydration at 10 minutes for all three *SNAP33 snap33-2 hpSNAP33* transgenic lines when compared to the *SNAP33 snap33-2* heterozygote (Fig. 1e). When comparing the effects of the different *SNARE* mutants, the greatest reduction in Col-0 hydration at 10 min post-pollination was observed on stigmas from the *vamp722-1 hpVAMP721* lines #1 and #2, and the *SNAP33 snap33-2 hpSNAP33* line #3 (Fig. 1c-e).

**Fig. 1.**
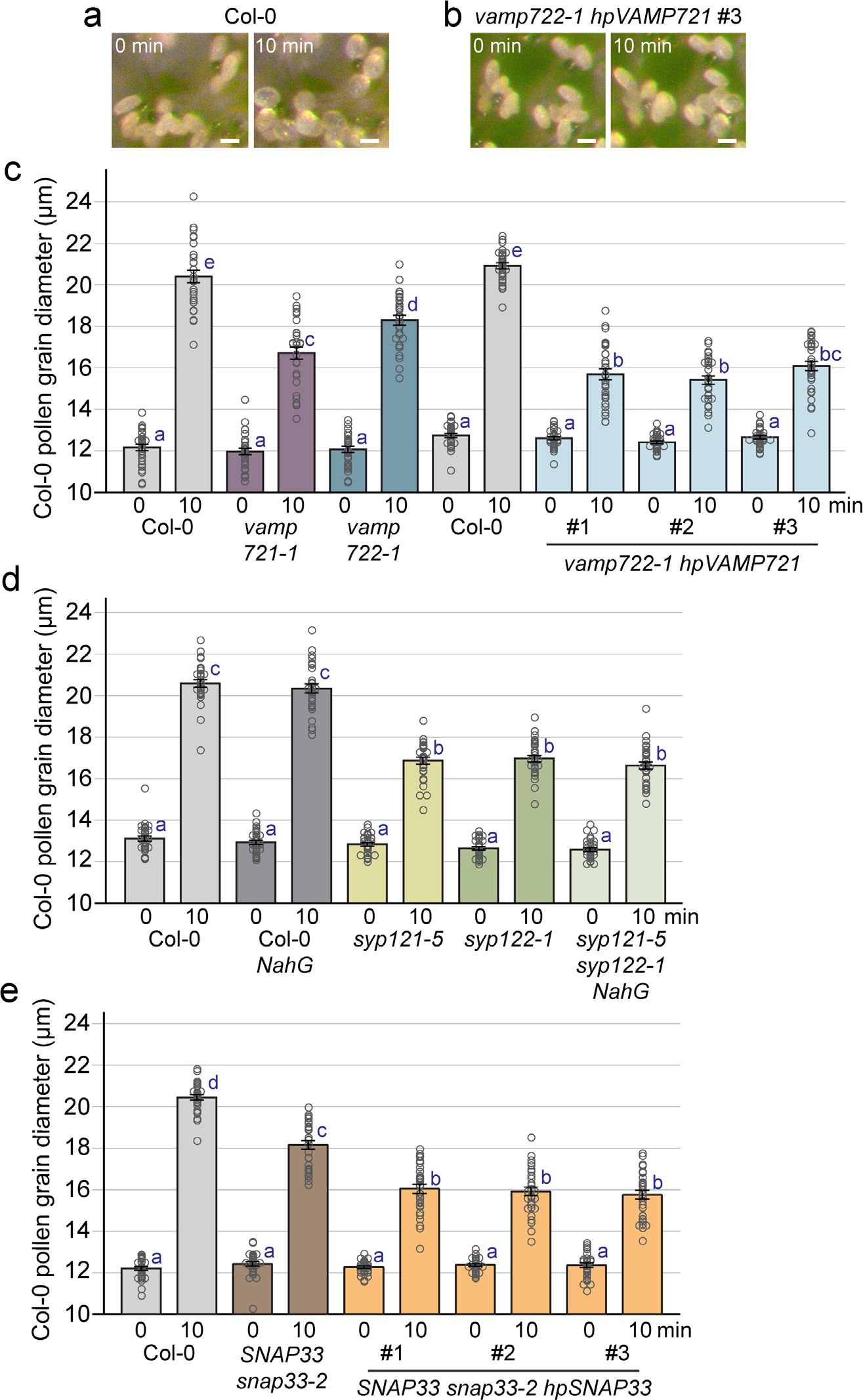
Col-0 pollen hydration assays on Col-0 and mutant SNARE stigmas. **a-b** Representative images of hydrating Col-0 pollen grains on stigmas from **a** Col-0 and **b** *vamp722-1 hpVAMP721* line #3. Scale bar = 20 µm. **c-e** Bar graphs displaying Col-0 pollen hydration on stigmas from **c** Col-0, the *vamp721-1* and *vamp722-1* single mutants, and three independent *vamp722-1 hpVAMP721* lines. **d** Col-0, Col-0 *NahG*, the *syp121-5* and *syp122-1* single mutants, and the *syp121-5 syp122-1 NahG* double mutant line. **e** Col-0, the *SNAP33 snap33-2* heterozygote, and three independent *SNAP33 snap33-2 hpSNAP33* lines. Col-0 pollen diameter was measured at 0 min and 10 min post-pollination. Data are shown as mean ± SE with all data points displayed. n = 30 pollen grains per line. Letters represent statistically significant groupings of P<0.05 based on a one-way ANOVA with a Tukey-HSD post-hoc test.

### Wildtype Col-0 pollen tube growth through *SNARE* mutant stigmas

Despite the reduced Col-0 pollen hydration at 10 min post-pollination on the *SNARE* mutant stigmas, the Col-0 pollen grains do hydrate sufficiently to germinate and form a pollen tube. To investigate Col-0 pollen tube growth through the mutant stigmas, aniline blue staining was performed at 24 hours post-pollination. Aniline blue stains callose which is deposited in the growing pollen tube, thus allowing for the visualization of the pollen tubes. Wildtype Col-0 stigmas can readily support pollen tube growth, and many pollen tubes are visible throughout the stigma and style and into the transmitting tract (Fig. 2a, b). Stigmas from the single *vamp721-1* and *vamp722*-1 mutants, and the three *vamp722-1 hpVAMP721* transgenic lines appeared to all support wildtype Col-0 pollen tube growth as well (Fig. 2c-l). These mutant pistils were lightly pollinated to more clearly observe the growing pollen tubes, but no differences were observed. More heavily Col-0-pollinated *vamp722-1 hpVAMP721* pistils stained at 2-hours and 24-hours post-pollination also displayed wild-type patterns of Col-0 pollen tube growth (Supplementary Fig. 6). Similarly, Col-0 pollen tubes grew normally through the stigmas from the single *syp121-5* and *syp122-1* mutants, and the double *syp121-5 syp122-1 NahG* mutant (Fig. 2m-r). Finally, wildtype Col-0 growth was observed in stigmas from the *SNAP33 snap33-2* heterozygote and the three *SNAP33 snap33-2 hpSNAP33* lines (Fig. 2s-z). The only potential phenotype we observed was that some Col-0 pollen tubes seemed to be more curled at the top of the *SNARE* mutant stigmas (e.g. Fig. 2x), however, this was not consistently seen with all pollen tubes. Overall, there was not a significant disruption in Col-0 pollen tube growth through the SNARE mutant stigmas. Together with the pollen hydration data, these data indicate that Col-0 pollen hydration may be delayed on the SNARE mutant stigmas, but this delay is eventually overcome, and Col-0 pollen germination and pollen tube growth occurs.

**Fig. 2.**
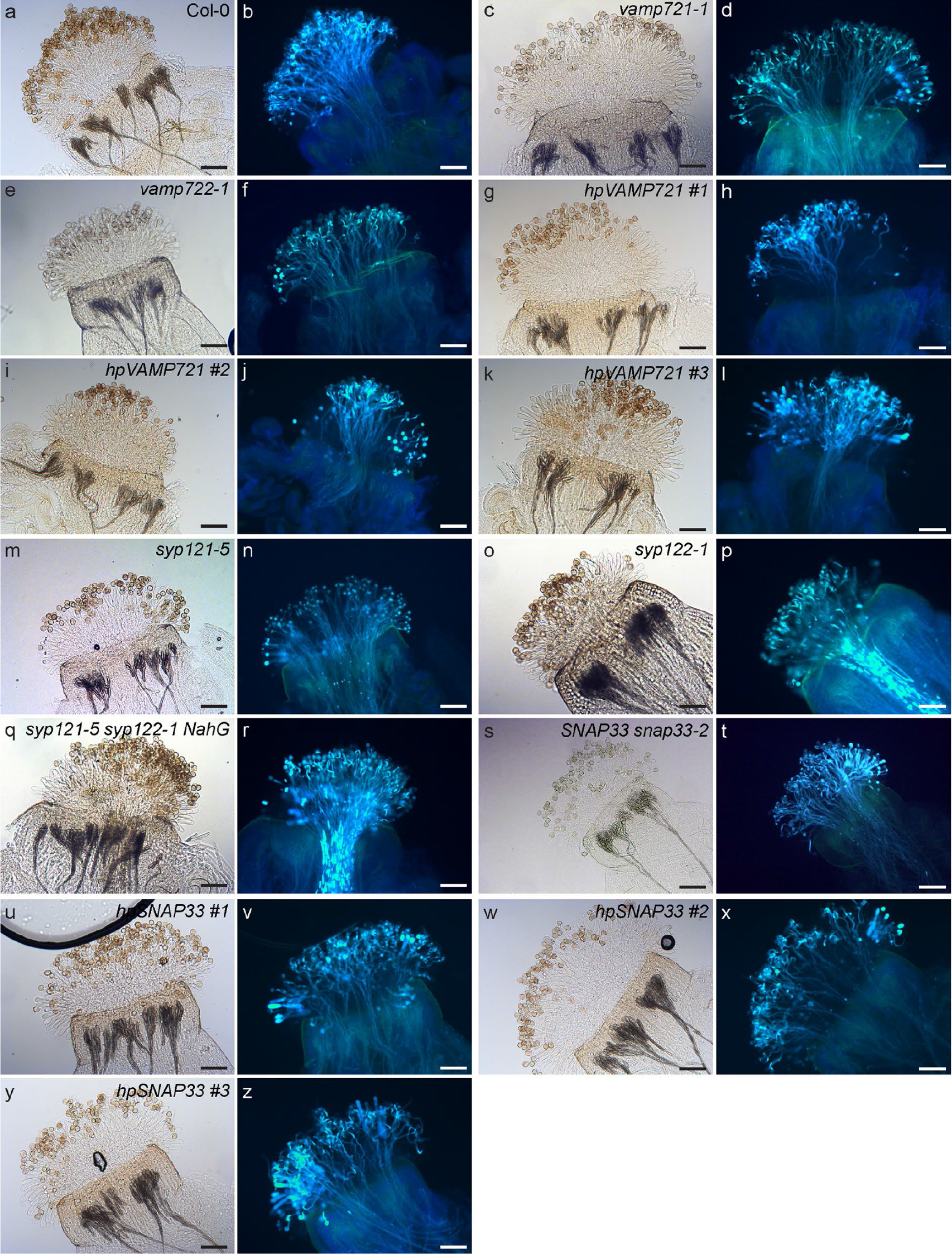
Representative images of aniline blue-stained Col-0 and mutant SNARE pistils pollinated with Col-0 pollen. **a-b** Col-0 pistil. **c-l** Pistils from *vamp721-1, vamp722-1*, and three independent *vamp722-1 hpVAMP721* lines. **m-r** Pistils from *syp121-5, syp122-1*, and *syp121-5 syp122-1 NahG* plants. **s-z** Pistils from *SNAP33 snap33-2*, and three independent *SNAP33 snap33-2 hpSNAP33* lines. All pistils were pollinated with Col-0 pollen for 24 hours and then stained with aniline blue. Representative brightfield and aniline blue images of each pistil are shown. The mutant pistils were lightly pollinated to better view the growing pollen tubes. Scale bar = 100 µm.

### Seed set following wildtype Col-0 pollinations on *SNARE* mutant pistils

While the focus of this study was on investigating the role of *SNAREs* in the Arabidopsis stigma with pollinations, we noticed that some mutants appeared to produce fewer seeds. To examine seed production, pistils from Col-0 and the different *SNARE* mutants were hand pollinated with Col-0 pollen, and the resulting siliques were harvested, and their seeds counted. While the siliques from the *vamp721-1* and *vamp722-1* mutants only had a slight decrease in seed yield compared to Col-0, the *vamp722-1 hpVAMP721* line #1 displayed a much larger decrease in seed set (Fig. 3a). The reduced seed set was unique to this *vamp722-1 hpVAMP721* line as the other two lines displayed wildtype seed set levels (Fig 3a). For line #1, it appeared to be caused by embryo lethality as the number of ovules per silique was similar to Col-0 (Supplementary Fig. 7). Siliques from the single *syp121-5* and *syp122-1* mutants, and the double *syp121-5 syp122-1 NahG* mutant all produced large amounts of seeds that were only slightly less than Col-0 (Fig. 3b). Similarly, the *SNAP33 snap33-2* heterozygote produced seeds/silique values that were similar to Col-0 (Fig. 3c). However, siliques from all three *SNAP33/snap33-2hpSNAP33* lines had significantly decreased seed yields compare to Col-0 (Fig. 3c). Given the consistent phenotype across all three *SNAP33 snap33-2 hpSNAP33* lines, we investigated if this reduced seed set might be linked to the number of ovules in each pistil. There was a significant decrease in the number of ovules per silique compared to Col-0 which would contribute to the decreased seed set observed (Supplementary Fig. 8). While the stigma-specific SLR1 promoter was used to drive the expression of *hpSNAP33*, it may be that the small RNAs generated from this construct were mobile and affected ovule development in these transgenic lines as well.

**Fig. 3.**
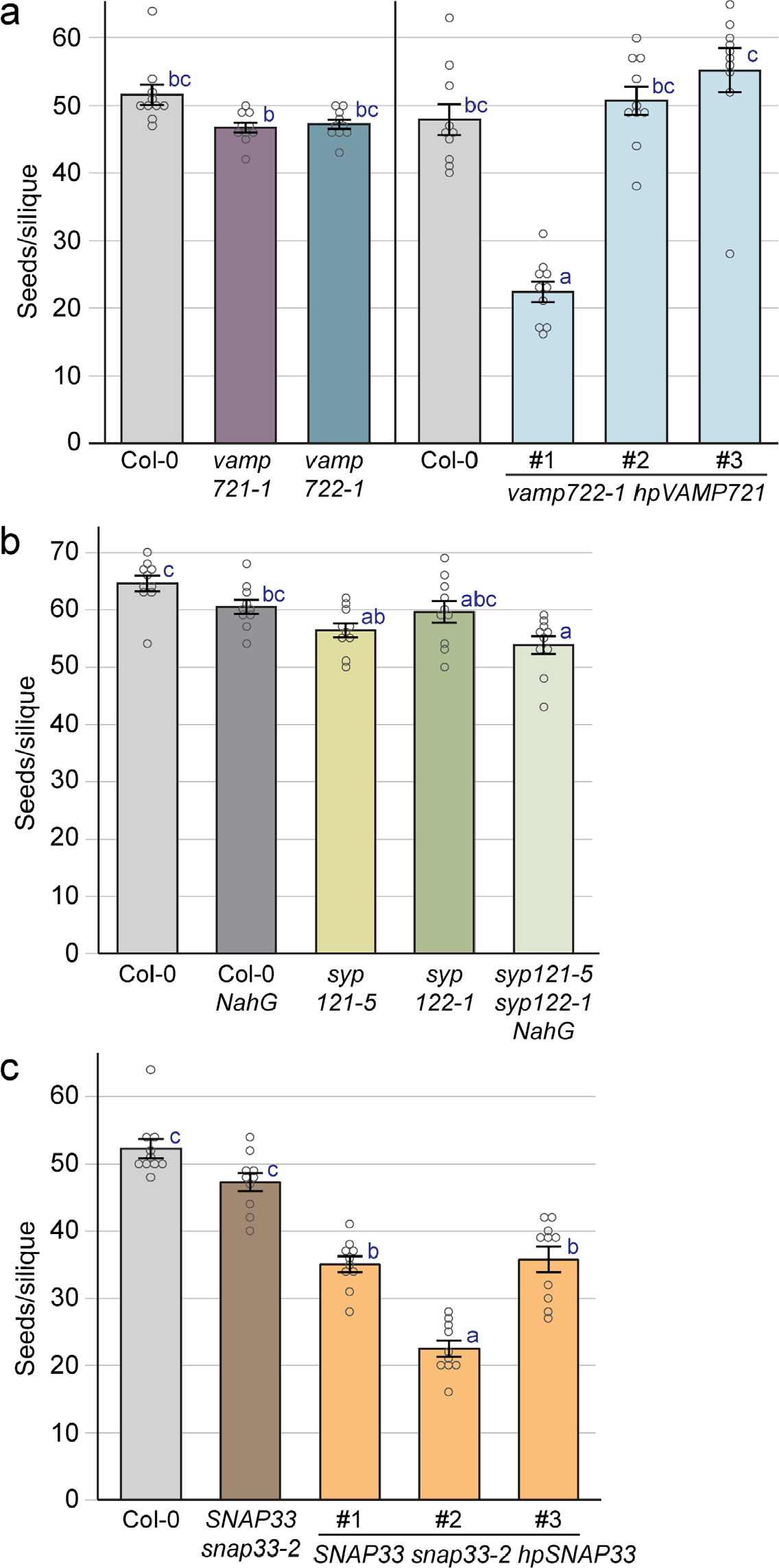
Seed production following Col-0 pollinated Col-0 and mutant SNARE pistils. Seed set following Col-0 pollination on pistils from **a** Col-0, the *vamp721-1* and *vamp722-1* single mutants, and three independent *vamp722-1 hpVAMP721* lines. **b** Col-0, Col-0 *NahG*, the *syp121-5* and *syp122-1* single mutants, and the *syp121-5 syp122-1 NahG* double mutant line. **c** Col-0, the *SNAP33 snap33-2* heterozygote, and three independent *SNAP33 snap33-2 hpSNAP33* lines Data are shown as mean ± SE with all data points displayed. n = 10 siliques per line. Letters represent statistically significant groupings of P<0.05 based on a one-way ANOVA with a Tukey-HSD post-hoc test.

## Discussion

In this study, knockout/knockdown lines for five Arabidopsis *SNARE* genes (*VAMP721, VAMP722, SYP121, SYP122* and *SNAP33*) were analyzed for pollen-stigma responses. These five *SNARE* genes were selected because, in addition to being expressed in the stigma (Supplementary Fig. 1), the proteins have been shown to form a protein complex to act as a membrane fusion complex (El Kasmi et al. 2013; Kwon et al. 2008). Different SNARE proteins contribute α-helical SNARE domains to form a four-helix bundle in the SNARE fusion complex. The vesicle R-SNARE (VAMP721/722) and plasma membrane Qa-SNARE (SYP121/122) each contribute one SNARE domain, and the plasma membrane Qbc-SNARE (SNAP33) contributes two SNARE domains to the four-helix bundle. The assembly of this complex brings the vesicle membrane and the plasma membrane together for fusion (Lipka et al. 2007; Luo et al. 2022; Shi et al. 2023). The data presented here supports a role for these five Arabidopsis *SNAREs* in the stigma to promote pollen hydration (Fig. 1). Stigmas from single *SNARE* mutants displayed significantly reduced wildtype Col-0 pollen hydration levels at 10 min post-pollination. Col-0 pollen hydration on stigmas from a *syp121-5 syp122-1* double mutant expressing *NahG* (to reduce SA levels) were similar to those of the single mutants. However, when the *vamp722-1* and *SNAP33 snap33-2* mutants were combined with their respective stigma-expressed hairpin RNAi constructs, *hpVAMP721* and *hpSNAP33*, there was a further reduction in Col-0 pollen hydration on the mutant stigmas. Thus, these data support a role for these five SNAREs in the stigma to support pollen hydration, presumably by facilitating vesicle fusion during pollen-induced stigmatic secretion.

Despite the reduced Col-0 pollen hydration at 10 min post-pollination on the mutant *SNARE* stigmas, the Col-0 pollen successfully germinated, and abundant pollen tube growth through the SNARE mutant stigmas/styles was observed (Fig. 2; Supplementary Fig. 6). Previously, using knockdown/knockout mutants for the eight exocyst subunit genes, we generally observed a stronger reduction in the ability of the mutant Arabidopsis stigmas to support the early pollen-stigma interactions compared to what was observed in this study for the five *SNARE* genes (Safavian et al. 2014; Safavian et al. 2015; Samuel et al. 2009). Due to the many pleiotropic effects caused by knocking out SNARE complex genes, it is challenging to get a good knockout of redundant *SNAREs* while avoiding overall plant growth defects, though the same would hold true for the exocyst subunit genes. There are a couple of possible explanations as to why we observed milder phenotypes in this study. For both *SNAP33* and *VAMP721/722*, the observed phenotypes in this study may simply be due to inefficient silencing by their respective hpRNAi constructs. However, there are also other potential *SNARE* genes that could be functioning redundantly with the *SNAREs* selected for this study (Supplementary Fig. 1). For example, the SYP132 Qa-SNARE functions in the constitutive secretion pathway during growth along with VAMP721/722 (Yun et al. 2013). The *SYP132* gene is broadly expressed (Supplementary Fig. 1) and may function redundantly with *SYP121/122* which is presumably why *syp121-5 syp122-1 NahG* plants are wildtype in appearance. For *VAMP721/722*, other *VAMP72* members may also be involved based on expression in the stigma (e.g. *VAMP724*, Supplementary Fig. 1). However, for *SNAP33*, the other *SNAP* genes are not expressed in the stigma and so other redundant candidates are less clear. Interestingly in plants, a second type of SNARE fusion complex has been proposed to be made up of four SNARE proteins (R-SNARE, Qa-SNARE, Qc-SNARE, Qc-SNARE) each contributing an α-helical SNARE domain to the four helix bundle (Lipka et al. 2007; Luo et al. 2022; Shi et al. 2023). In cell plate formation, the Knolle/SYP111 Qa-SNARE and the SNAP33 Qbc-SNARE were found to form a complex with the VAMP721/722 R-SNAREs, but an alternate complex was also discovered consisting of the Knolle/SYP111 Qa-SNARE, the NPSN11 Qb-SNARE, the SYP71 Qc-SNARE, and the VAMP721/722 R-SNAREs (El Kasmi et al. 2013). Double mutant analyses of *SNAREs* in each complex (e.g. *snap33 npsn11* and *snap33 syp71*) resulted in stronger cytokinesis defects in the embryo implicating both complexes in cytokinesis (El Kasmi et al. 2013). Both SYP71 and NPSN homologues are expressed well in the Arabidopsis stigma (Supplementary Fig. 1) and could perhaps replace the SNAP33 function in the stigma in response to pollen.

In this study, we made the assumption that the exocyst and SNARE complexes functioned in a typical model where the exocyst complex is responsible for tethering vesicles at the plasma membrane prior to SNARE-mediated membrane fusion to release the vesicle cargo (Saeed et al. 2019). Consistent with this model, the exocyst subunit, EXO70A1, was found to interact with SYP121, SNAP33 and VAMP721/722 (Larson et al. 2020). However, a recent study has proposed that there are two parallel Rab GTPase-regulated pathways for vesicle secretion (Nielsen 2022; Pang et al. 2022). In the typical pathway, a Rab GTPase recruits the exocyst complex to tether the vesicle at the plasma membrane, followed by SNARE-mediated fusion. However, Pang et al. (2022) discovered a second pathway that is exocyst-independent where the RabA2a Rab GTPase directly interacts with SNARE proteins to promote the formation of the SNARE fusion complex; all of which highlights the increasing complexities being discovered for SNARE mediated membrane fusion in plants. Interestingly, we have previously uncovered that the self-incompatibility pathway in Brassicaceae species targets EXO70A1 in the stigmatic papillae to block pollen hydration (Abhinandan et al. 2022; Goring 2017; Goring et al. 2023; Samuel et al. 2009). By targeting this exocyst component, the SNAREs would not be able to follow up with vesicle fusion to the plasma membrane. This also implies that the second direct RabA2a-SNARE pathway may not be active to compensate and promote vesicle secretion for pollen hydration. Overall, our data implicates Arabidopsis SNAREs functioning in the stigma for promoting pollen hydration and provides insights into how the individual *VAMP721, VAMP722, SYP121, SYP122* and *SNAP33* genes contribute to this process. These findings also add further support to the centrality of the secretory system in the Arabidopsis stigmatic papillae following pollinations.

## Acknowledgements

We thank Tracy Fang, Judy Fang and Yihan Gong for technical assistance, and members of the Goring lab for critically reading this article. We are also very grateful to Dr. Chian Kwon (Dankook University) and the ABRC for providing T-DNA mutant seed stocks, and Dr. Keiko Yoshioka (University of Toronto) for providing the *NahG* transgenic seeds. SRM was supported by a Rustom H. Dastur Graduate Scholarship, and this research was supported by a grant from the Natural Sciences and Engineering Research Council of Canada to DRG.

