## Supplemental Figures and Tables for "The Arabidopsis *SNARE* complex genes regulate the early stages of pollen-stigma interactions"

| Name | Stigmatic tissue - 6th and 7th flowers | Ovules - 6th and 7th flowers | Carpel - 6th and 7th flowers (ovules & stigmas removed) | Opened anthers, with pollen | Anthers - mature flower (before opening) | Carpels - mature flower (before pollination) | Stamen filaments - mature flower | Petals - mature flower | Sepals - mature flower | Pedicel | Young anther | Young carpel | Young sepal | Flower 1 | Young seeds 1 |
|---|---|---|---|---|---|---|---|---|---|---|---|---|---|---|---|
| SYP111 | 136 | 2675 | 2786 | 126 | 49 | 3097 | 113 | 47 | 68 | 3156 | 828 | 4877 | 755 | 1328 | 1248 |
| SYP112 | 10 | 12 | 42 | 85 | 256 | 39 | 23 | 15 | 61 | 51 | 337 | 39 | 19 | 35 | 7 |
| SYP121 | 760 | 839 | 1172 | 1334 | 121 | 993 | 1128 | 1402 | 1033 | 757 | 659 | 1506 | 2144 | 1177 | 770 |
| SYP122 | 83 | 66 | 100 | 333 | 42 | 241 | 96 | 180 | 1748 | 89 | 72 | 91 | 1086 | 538 | 36 |
| SYP123 | 6 | 6 | 72 | 206 | 10 | 79 | 10 | 4 | 0.5 | 130 | 62 | 16 | 22 | 54 | 10 |
| SYP124 | 2 | 2 | 5 | 11037 | 15337 | 5 | 36 | 10 | 8 | 2 | 9 | 12 | 4 | 392 | 6 |
| SYP125 | 4 | 0 | 0 | 4381 | 9770 | 1 | 16 | 6 | 1 | 0 | 0 | 0 | 4 | 212 | 4 |
| SYP131 | 23 | 2 | 2 | 21980 | 22433 | 10 | 42 | 15 | 12 | 2 | 25 | 0 | 2 | 1020 | 8 |
| SYP132 | 1256 | 915 | 1107 | 529 | 937 | 1012 | 3675 | 3544 | 1059 | 1304 | 727 | 975 | 1130 | 996 | 961 |
| NPSN11 | 919 | 962 | 949 | 2933 | 5040 | 984 | 2412 | 1275 | 289 | 1354 | 337 | 1264 | 508 | 634 | 983 |
| NPSN12 | 410 | 448 | 560 | 787 | 798 | 516 | 1310 | 793 | 472 | 1063 | 298 | 554 | 484 | 464 | 617 |
| NPSN13 | 804 | 911 | 835 | 1748 | 2050 | 738 | 1716 | 1416 | 772 | 940 | 549 | 939 | 767 | 625 | 859 |
| SYP71 | 2129 | 1860 | 1492 | 847 | 659 | 1452 | 3999 | 2840 | 1902 | 1861 | 1529 | 1552 | 1458 | 1302 | 1807 |
| SYP72 | 11 | 0 | 1 | 6601 | 8998 | 3 | 14 | 13 | 17 | 5 | 127 | 0 | 3 | 332 | 1 |
| SYP73 | 1 | 18 | 22 | 98 | 119 | 15 | 6 | 10 | 2 | 10 | 282 | 6 | 2 | 33 | 11 |
| SNAP29 | 5 | 18 | 5 | 11 | 9 | 10 | 4 | 5 | 6 | 5 | 43 | 10 | 1 | 15 | 20 |
| SNAP30 | 1 | 3 | 1 | 4232 | 9247 | 3 | 11 | 1 | 3 | 0 | 2 | 0 | 0.5 | 146 | 3 |
| SNAP33 | 1496 | 1156 | 1092 | 1250 | 594 | 1162 | 1202 | 1264 | 1892 | 1110 | 1107 | 1142 | 1969 | 1740 | 975 |
| VAMP721 | 3313 | 2459 | 2404 | 1258 | 913 | 2491 | 4054 | 3208 | 1936 | 2912 | 1777 | 2340 | 2085 | 2156 | 2445 |
| VAMP722 | 3162 | 1343 | 1148 | 2293 | 1486 | 1486 | 3665 | 2368 | 1995 | 1503 | 1220 | 1136 | 1450 | 1501 | 1489 |
| VAMP724 | 855 | 1125 | 1263 | 276 | 194 | 1152 | 1123 | 1537 | 917 | 1188 | 1303 | 1837 | 1096 | 1013 | 1060 |
| VAMP725 | 62 | 147 | 65 | 295 | 4148 | 84 | 85 | 84 | 42 | 60 | 155 | 54 | 62 | 63 | 64 |
| VAMP726 | 11 | 189 | 170 | 1304 | 4399 | 160 | 14 | 31 | 17 | 206 | 123 | 247 | 67 | 130 | 81 |

| Scale |
|---|
| 0 |
| 500 |
| 1000 |
| 1500 |
| 2000 |
| 2500 |
| 3000 |
| 3500 |
| 4000 |
| 4500 |
| >5000 |

| Localization* | AGI* | Type* | Name* | Stage 13 Receptive stigmatic papillae |
|---|---|---|---|---|
| CP/TGN/MVB | AT1G08560 | Qa | SYP111 | |
| PM | AT2G18260 | Qa | SYP112 | 5 |
| PM | AT3G11820 | Qa | SYP121 | 36 |
| PM | AT3G52400 | Qa | SYP122 | 16 |
| PM | AT4G03330 | Qa | SYP123 | |
| PM | AT1G61290 | Qa | SYP124 | 1 |
| PM | AT1G11250 | Qa | SYP125 | |
| PM | AT3G03800 | Qa | SYP131 | 2 |
| PM/CP | AT5G08080 | Qa | SYP132 | 68 |
| CP/TGN/PM | AT2G35190 | Qb | NPSN11 | 46 |
| TGN/PM | AT1G48240 | Qb | NPSN12 | 33 |
| TGN/PM | AT3G17440 | Qb | NPSN13 | 30 |
| CP/PM/E/ER | AT3G09740 | Qc | SYP71 | 91 |
| ER/PM | AT3G45280 | Qc | SYP72 | 1 |
| ER/PM | AT3G61450 | Qc | SYP73 | 1 |
| PM | AT5G07880 | Qbc | SNAP29 | |
| PM | AT1G13890 | Qbc | SNAP30 | |
| CP/PM/E | AT5G61210 | Qbc | SNAP33 | 106 |
| PM/TGN/EE/CP | AT1G04750 | R | VAMP721 | 118 |
| PM/TGN/EE/CP | AT2G33120 | R | VAMP722 | 175 |
| TGN/PM | AT4G15780 | R | VAMP724 | 21 |
| TGN/PM | AT2G32670 | R | VAMP725 | 4 |
| TGN/PM | AT1G04760 | R | VAMP726 | 1 |

| Scale |
|---|
| 0 |
| 20 |
| 40 |
| 60 |
| 80 |
| 100 |
| 120 |
| 140 |
| 160 |
| 180 |
| >200 |

**Supplementary Fig. 1** Expression profiles of plasma membrane SNARE genes in floral tissue RNA-Seq datasets.

* Retrieved from Table 3 from Luo et al. 2022. SNAREs Regulate Vesicle Trafficking During Root Growth and Development. Front Plant Sci. 13:853251. https://doi.org/10.3389/fpls.2022.853251

Stigmatic papillae RNA-Seq data: Gao et al. 2018. KIRA1 and ORESARA1 terminate flower receptivity by promoting cell death in the stigma of Arabidopsis. Nat Plants 4:365-375. https://doi.org/10.1038/s41477-018-0160-7

Floral tissue RNA-Seq data: TMM-normalized raw values retrieved from TRAVA http://travadb.org/ Klepikova et al. 2016. A high resolution map of the Arabidopsis thaliana developmental transcriptome based on RNA-seq profiling. Plant J 88:1058-1070.

Heatmaps generated with BAR HeatMapper Plus Tool. http://bar.utoronto.ca/
Toufighi et al. 2005. The Botany Array Resource: e-Northerns, Expression Angling, and Promoter analyses. Plant J 43:153-163.

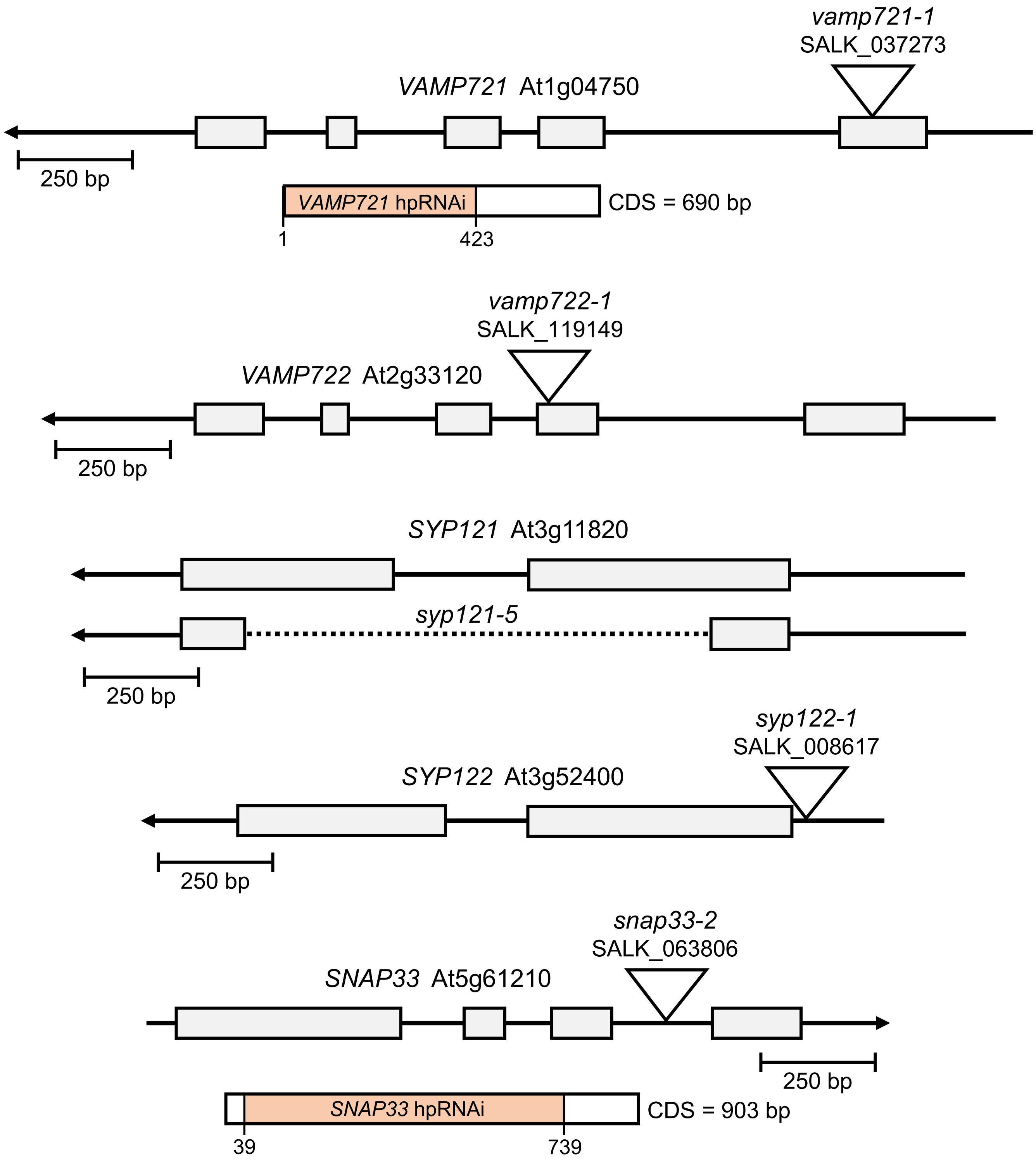

**Supplementary Fig. 2** SNARE mutants used in this study.
Previously characterized T-DNA mutants in the Col-0 accession used in this study were *vamp721-1* (SALK_037273; Kwon et al. 2008), *vamp722-1* (SALK_119149; Kwon et al. 2008), *syp122-1* (SALK_008617; Assaad et al. 2004), and *snap33-2* (SALK_063806; Henchiri 2021). The *syp121-5* mutant in the Col-0 background was created using CRISR/Cas9 (see Supplementary Fig. 4). The cDNA fragments used in the hpRNAi vectors targeting *VAMP721* and *SNAP33* are shown in orange. The *snap33-2* mutant was used in this study as the previously published *snap33-1* mutant was in Ws accession (Heese et al. 2001).

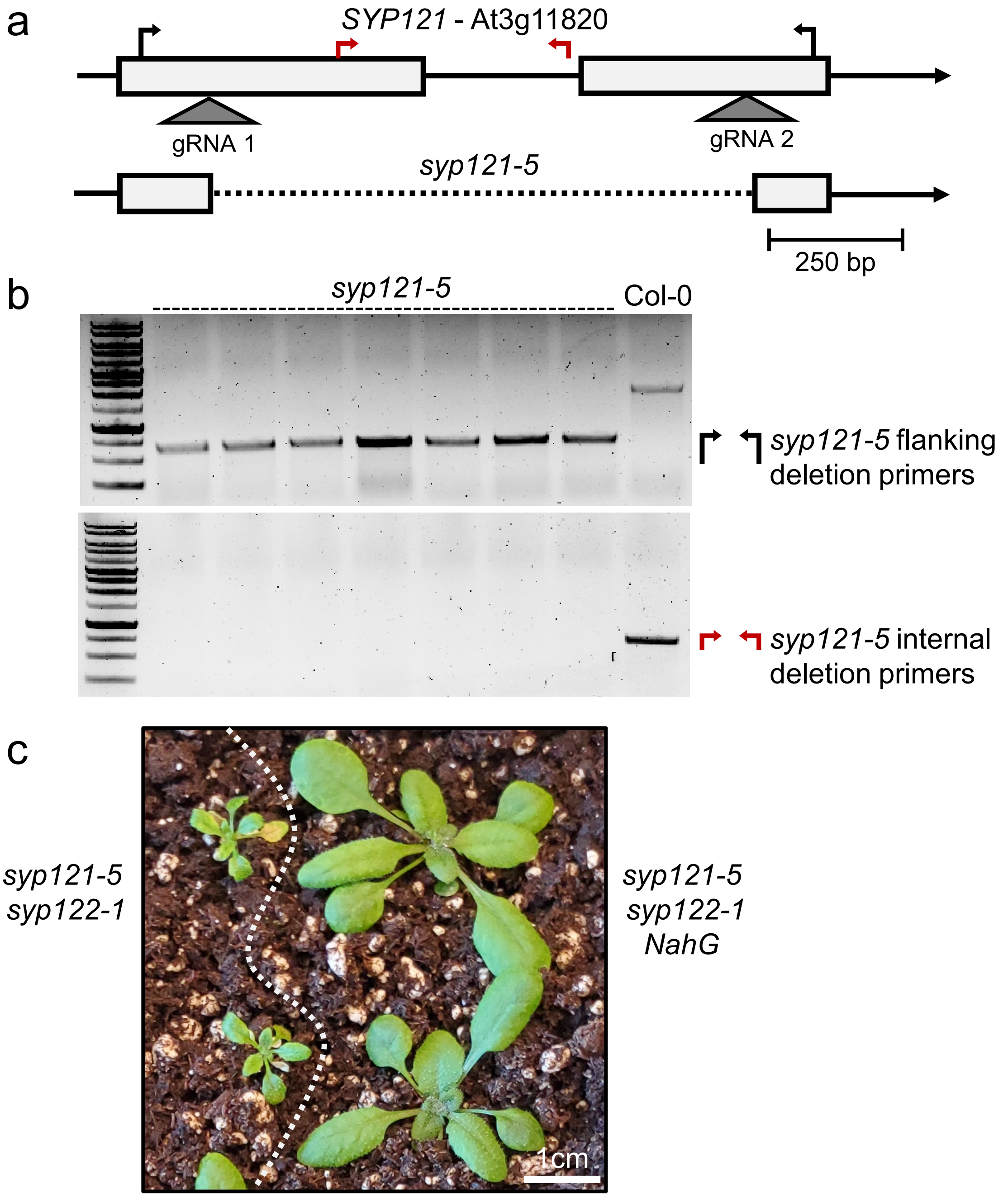

**Supplementary Fig. 3** Generation of the *syp121-5* deletion mutation.
**a** Schematic of the *SYP121* gene showing the location of the two gRNAs targeting exon 1 and exon 2 for the CRISPR/Cas9-mediated deletion. Genotyping primers are also shown above the gene. The *syp121-5* mutant was generated to be easier to track in crosses since the previously characterized syp121-1 mutant was an EMS mutant (ref).
**b** The *syp121-5* deletion was detected using primers flanking the gRNA cut sites (**a**, black). A smaller band is seen for the *syp121-5* mutant plants compared to Col-0 due to the internal deletion shown in (**a**). The $T_3$ progeny in lanes 1 to 7 are homozygous for the *syp121-5* mutation as no bands are seen using primers in the internal deleted region (**a**, red).
**c** The *syp121-5 syp122-1* double mutant displays the same seedling growth defects as previously characterized in the *syp121-1 syp122-1* double mutant The *NahG* transgene was crossed in to rescue these growth defects in the *syp121-5 syp122-1* double mutant resulting in wild-type growth and development (Zhang et al. 2008).

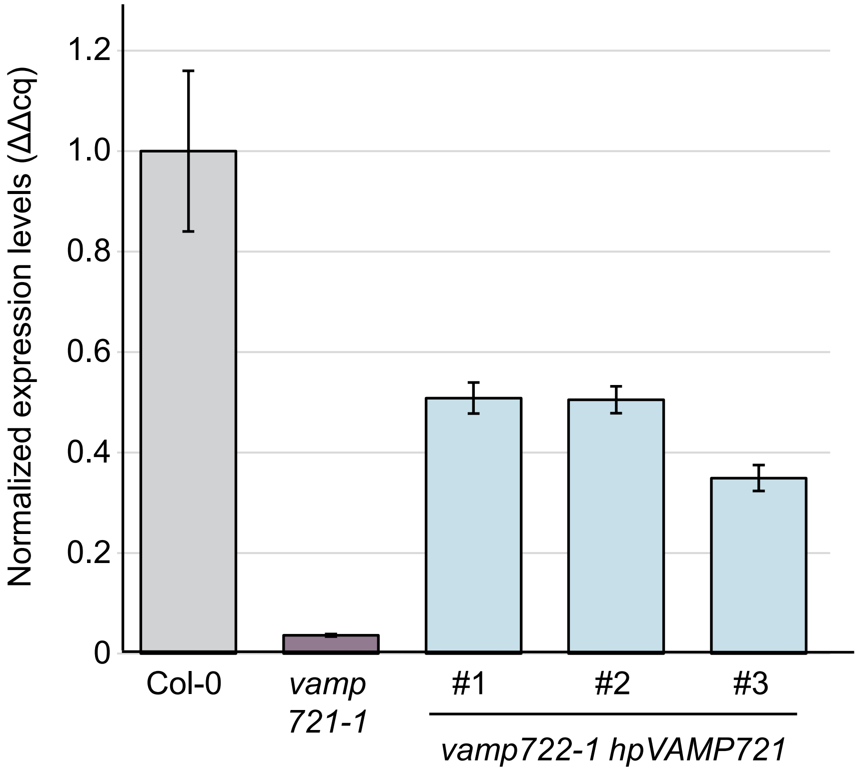

**Supplementary Fig. 4** RT-qPCR analysis of *VAMP721* expression.
RNA was extracted from the top-half of pistils collected from Stage 12 flower buds from Col-0, *vamp721-1*, and three different *hpVAMP721* lines in the *vamp722-1* mutant background. The addition of the *hpVAMP721* RNAi silencing construct resulted in a reduction of *VAMP721* expression.
*VAMP721* expression was normalized against *TUB6* and *ACT7* and plotted relative to Col-0.

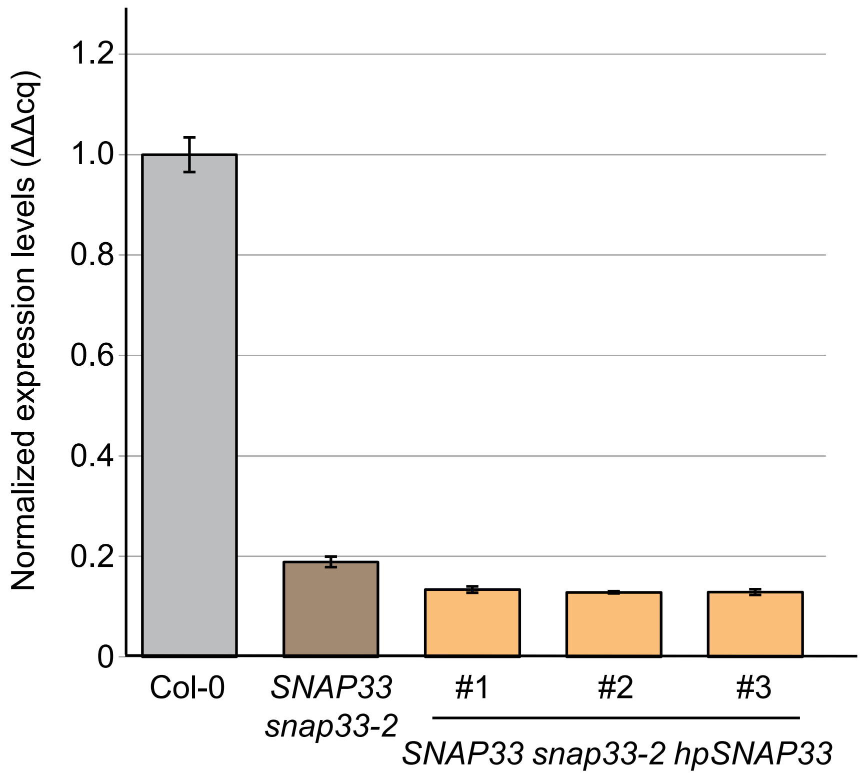

**Supplementary Fig. 5** RT-qPCR analysis of *SNAP33* expression.
RNA was extracted from the top-half of pistils collected from Stage 12 flower buds from Col-0, the *SNAP33/snap33-2* heterozygote, and three different *hpSNAP33* lines in the *SNAP33/snap33-2* background. The addition of the *hpSNAP33* RNAi silencing construct resulted in an even greater reduction of *SNAP33* expression. *SNAP33* expression was normalized against *TUB6* and *ACT7* and plotted relative to Col-0.

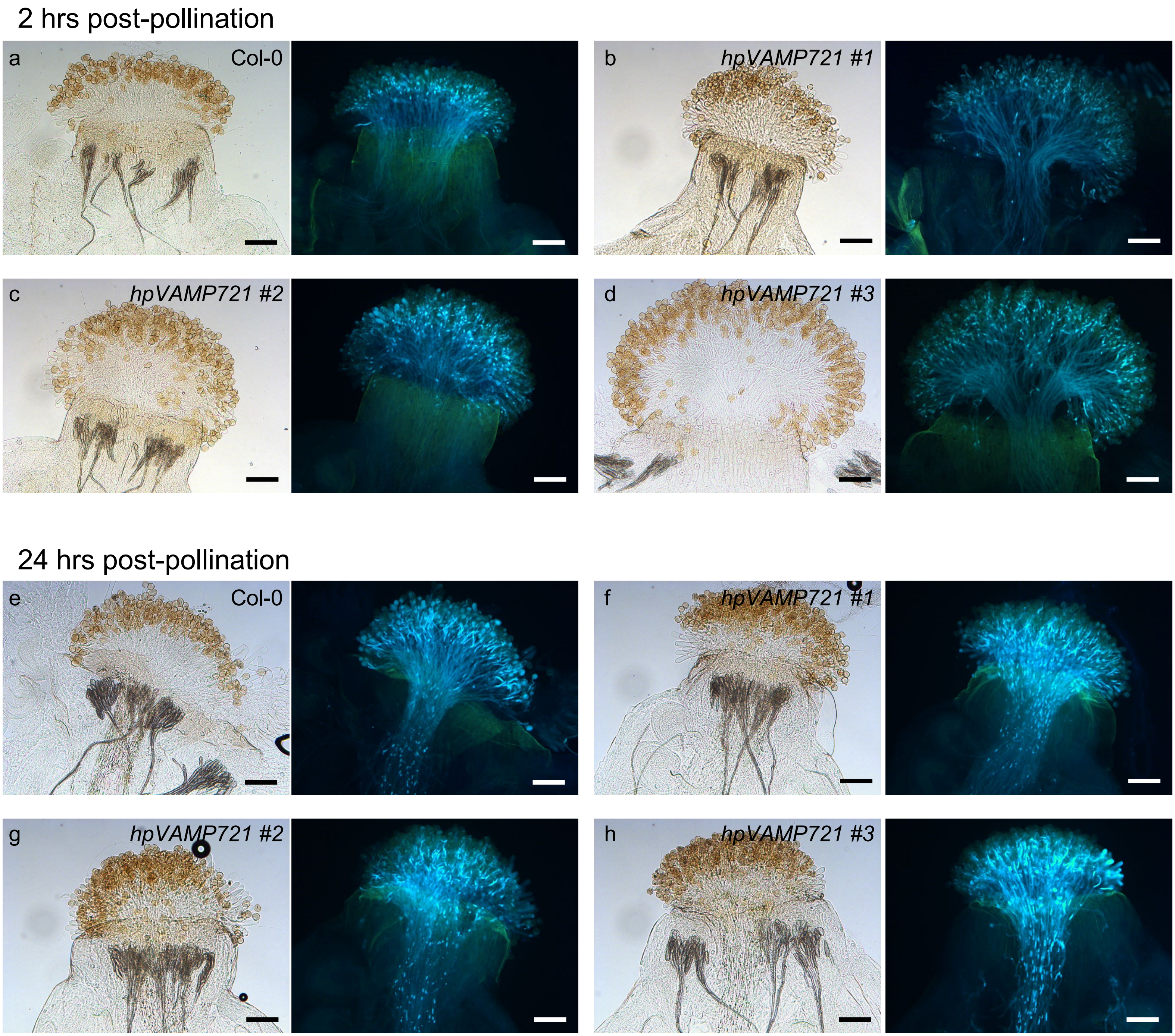

**Supplementary Fig. 6** Representative images of aniline blue-stained Col-0 and *vamp722-1 hpVAMP721* pistils pollinated with Col-0 pollen at 2 and 24 hrs post-pollination.
**a-d** 2 hrs post-pollination for pistils from Col-0 (**a**) and three independent *vamp722-1 hpVAMP721* lines (**b-d**).
**e-h** 24 hrs post-pollination for pistils from Col-0 (**e**) and three independent *vamp722-1 hpVAMP721* lines (**f-h**).
All pistils were pollinated with Col-0 pollen and then stained with aniline blue. Representative brightfield and aniline blue images of each pistil are shown. Scale bar = 100 µm.

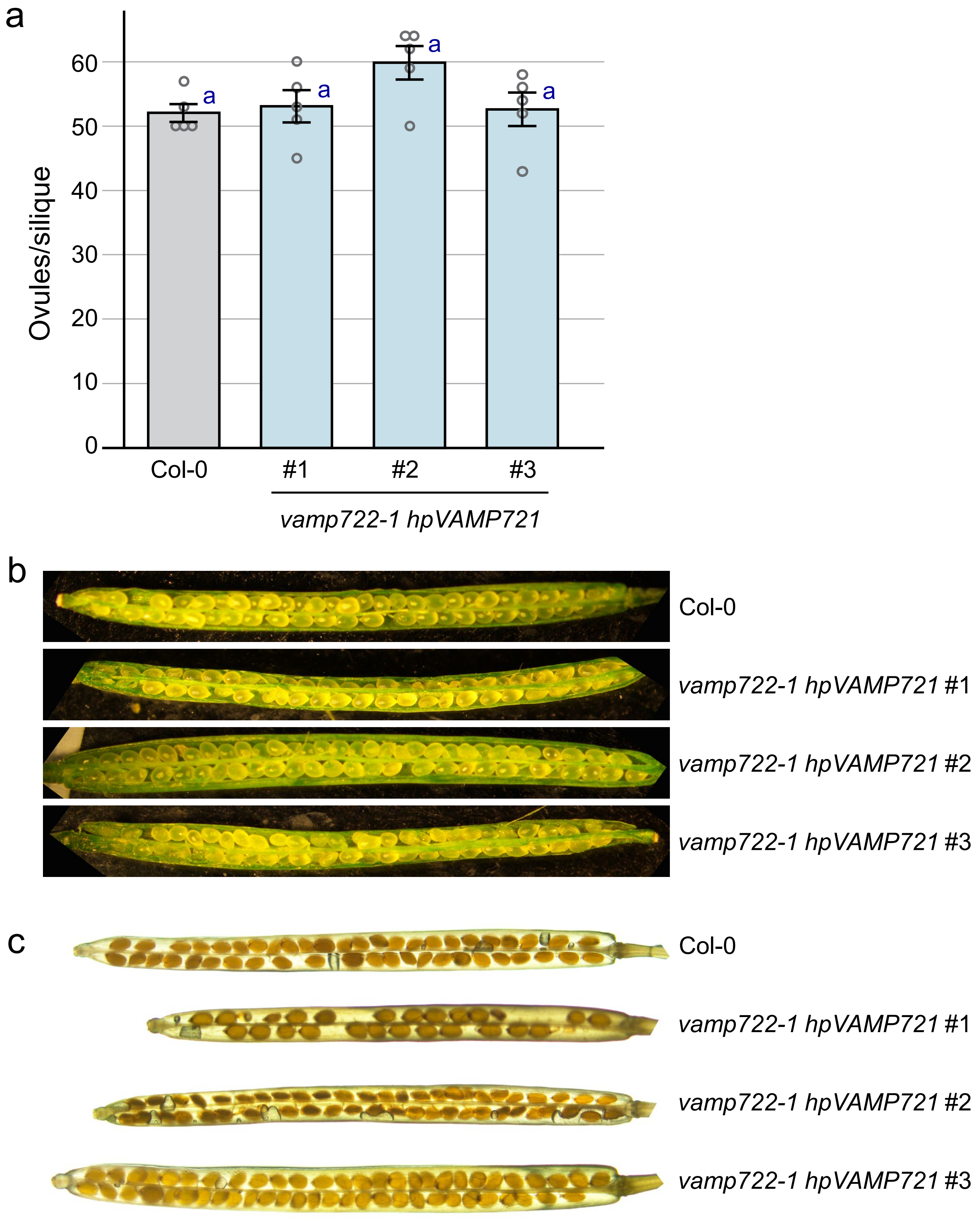

**Supplementary Fig. 7** Ovule counts for the *vamp722-1 hpVAMP721* lines.
**a** Ovules were counted from dissected siliques as described in Yuan and Kessler (2019). The *vamp722-1 hpVAMP721* lines did not show any significant differences in the number of ovules/siliques compared to Col-0. n= 5siliques per line, $P<0.05$ (One-way ANOVA with Tukey-HSD post-hoc test).
**b** Representative images of ovules in dissected siliques from Col-0 and the *vamp722-1 hpVAMP721* lines.
**c** Representative images of seeds in dissected mature siliques from Col-0 and the *vamp722-1 hpVAMP721* lines. Siliques from *vamp722-1 hpVAMP721* #1 had a reduced number of fully developed seeds (Figure 3). While a stigma-specific promoter was used to drive the expression of the *hpVAMP721* RNAi silencing construct, the small RNAs may have mobile affecting seed development in *vamp722-1 hpVAMP721* #1.

a

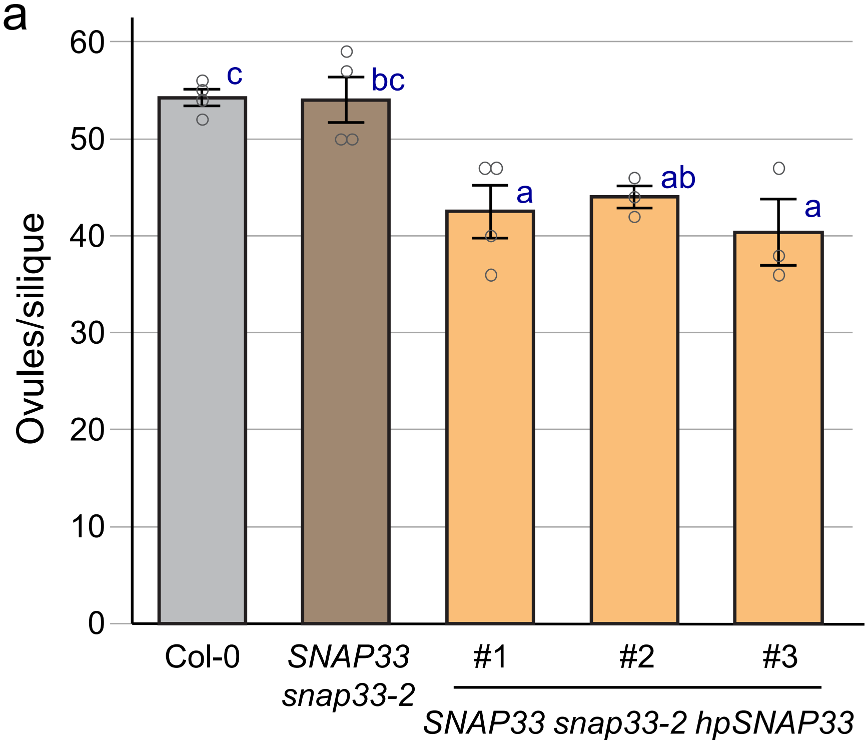

b

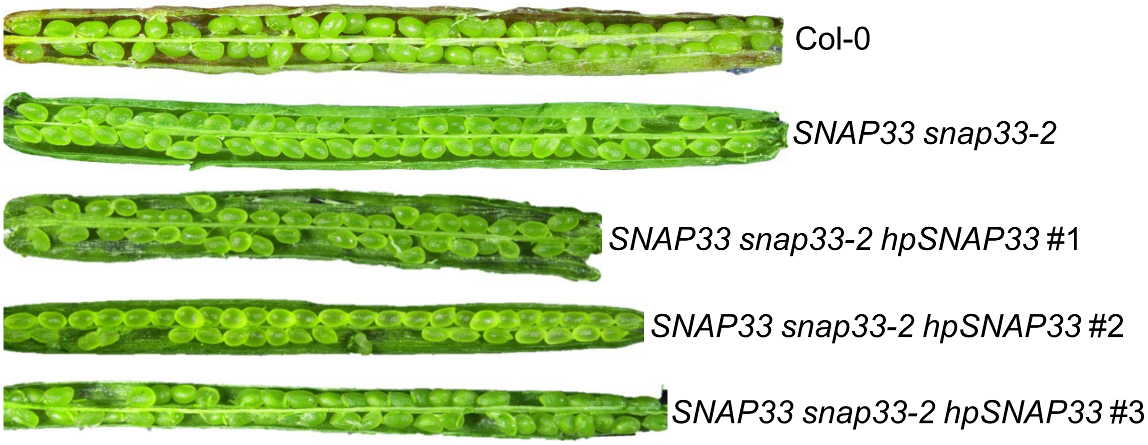

**Supplementary Fig. 8** Ovule counts for the *SNAP33 snap33-2 hpSNAP33* lines.
**a** Ovules were counted from dissected siliques as described in Yuan and Kessler (2019). The *SNAP33 snap33-2 hpSNAP33* lines showed a decrease in the number of ovules per silique. While a stigma-specific promoter was used to drive the expression of the *hpSNAP33* RNAi silencing construct, the small RNAs may have mobile affecting ovule development as well. n= 3-4 siliques per line, $P<0.05$ (One-way ANOVA with Tukey-HSD post-hoc test).
**b** Representative images of dissected siliques from Col-0 and each mutant line.

**Supplementary Table 1.** Primers used in this study.

| | | | |
|---|---|---|---|
| **CRISPR/Cas9 Primers Used** | | | |
| **Name** | **AGI** | | **Sequence** |
| *syp121-5* target site 1 with BsaI and gRNA scaffold | AT3G11820 | | ATATATGGTCTCGATTGAGTTCAGATGGCGAATCCCGGTT TTAGAGCTAGAAATAGCAAGTTAAAAT |
| *syp121-5* target site 2 with BsaI and gRNA scaffold | AT3G11820 | | ATTATTGGTCTCTAAACATTTGGTTGACTTCCTCCAGCAAT CTCTTAGTCGACTCTACCAATA |
| *syp121-5* deletion FP | AT3G11820 | | CTTACACGTACGGGACAAGG |
| *syp121-5* deletion RP | AT3G11820 | | GGGAGATTAACAACAAACCAGC |
| *syp121-5* internal FP | AT3G11820 | | GGACCTCTGTCCTCAATGGT |
| *syp121-5* internal RP | AT3G11820 | | CTGGTGAAGCTCCCTGAGAT |
| **Genotyping** | | | |
| **Name** | **AGI** | **T-DNA ID** | **Primer Sequences** |
| *Lb1.3* | | | ATTTTGCCGATTTCGGAAC |
| *snap33-2* | AT5G61210 | SALK_063806 | GATAAGCATCAGCTGATTCGG |
| | | | TTTTGGTTTCTGCAGGAAGG |
| *vamp721-1* | AT1G04750 | SALK_037273 | TGTCCTTGTGAACAACAGGG |
| | | | AGATGGAAATCCGGAGTGTG |
| *vamp722-2* | AT2G33120 | SALK_119149 | ACATCAATCGCTGCTCAGTG |
| | | | ACGGTCAAGAACCTGCAAAC |
| *syp122-1* | AT3G52400 | SALK_008617 | GATGAGACGCTCGAGGGTAG |
| | | | TGGTTTCCCTTCAGCAATTC |
| *NahG* | N/A | | TCCAGGTACAGCTGTTCGAG |
| | | | AAAGGTGAGGATATGGCCGT |
| **Cloning** | | | |
| **Name** | **AGI** | **CDS Region** | **Primer Sequence** |
| SNAP33 hpFragment attB1 | AT5G61210 | 39-739 bp | GGGGACAAGTTTGTACAAAAAAGCAGGCTTAATGCATAACT CAGTCGACCTCAAGT |
| SNAP33 hpFragment attB2 | | | GGGGACCACTTTGTACAAGAAAGCTGGGTGCTGAAAGCCC ATCGTCTTGCTT |
| VAMP721 hpFragment attB1 | AT1G04750 | 1-423 bp | GGGGACAAGTTTGTACAAAAAAGCAGGCTTAATGATGGCG CAACAATCGTTGATC |
| VAMP721 hpFragment attB2 | | | GGGGACCACTTTGTACAAGAAAGCTGGGTGACACTTAAAC CCATGGCAAAC |
| pOA-RQ-HPgatewayCassette SacI fragment | N/A | | ggtggtGAGCTCATGGCGCAACAATCGTTGATC |
| pOA-RQ-HPgatewayCassette ClaI fragment | N/A | | ggtggtATCGATGATCAACGATTGTTGCGCCAT |
| **qPCR** | | | |
| **Name** | **AGI** | **Orientation** | **Primer Sequence** |
| ACT7 | At5g09810 | FP | GGTGAAGGCTGGTTTTGCTG |
| ACT7 | At5g09810 | RP | CCATGACACCAGTGTGCCTA |
| UBQ5 | AT3G62250 | FP | CAACATCCAGAAGGAATCGAC |
| UBQ5 | AT3G62250 | RP | TCTTCGGCTTGGTGTAAGTCT |
| TUB6 | AT5G12250 | FP | TGCGACTGTCTTCAAGGTTTCC |
| TUB6 | AT5G12250 | RP | AGATCACCAAAGCTAGGAGTG |
| SNAP33 | AT5G61210 | FP | TCTCAGTCGTGGTGAGAAAC |
| SNAP33 | AT5G61210 | RP | CGTTGGTGAGTCATCTCTGG |
| VAMP721 | AT1G04750 | FP | GATGGATTCACCTATTGTGT |
| VAMP721 | AT1G04750 | RP | AGCCTTTCCACCACCATATC |
